# Uncovering the Binding Mode of *γ*-Secretase Inhibitors

**DOI:** 10.1101/611194

**Authors:** M. Hitzenberger, M. Zacharias

## Abstract

Knowledge of how transition state inhibitors bind to *γ*-secretase is of major importance for the design of new Alzheimer’s disease therapies. Based on the known structure of *γ*-secretase in complex with a fragment of the amyloid precursor protein we have generated a structural model of *γ*-secretase in complex with the effective L-685,458 transition state inhibitor. The predicted binding mode is in excellent agreement with experimental data, mimicking all enzyme-substrate interactions at the active site and forming the relevant transition state geometry with the active site aspartate residues. In addition, we found that the stability of the complex is very likely also sensitive to the pH value. Comparative simulations on the binding of L-685,458 and the epimer L682,679 allowed us to explain the strongly reduced affinity of the epimer for *γ*-secretase. The structural model could form a valuable basis for the design of new or modified *γ*-secretase inhibitors.

## Article

The intra-membrane cleaving hetero-tetramer *γ*-secretase (GSEC) processes the C-terminal fragment of the amyloid precursor protein (APP), C99.^1–6^ Since some of the resulting cleavage products are strongly linked to Alzheimer’s disease (AD), GSEC is at the focus of many drug-design research efforts.^7–11^ Ultimately, one wishes to control or modulate C99 processing to prevent the progression of, or even cure, AD.^12–17^ A large number of experimental^4, 7–9, 18–24^ as well as theoretical^25–34^ studies have been undertaken to investigate and characterize the structure, biology and chemistry of GSEC and its substrates. Not long ago, the structures of GSEC (inactive mutation) in complex with Notch and C83 (a shortened APP fragment) have been solved.^23, 24^ Also recently, the binding mode of the pseudo-inhibitor DAPT has been ascertained by computational methods.^35^ However, a detailed structure or molecular model of how an effective transition state inhibitor binds to presenilin (PS), the catalytically active domain of GSEC, is still missing. Such a structure could be very valuable for future modulator or inhibitor design strategies.

Based on the recent GSEC-substrate structures,^23, 24^ we performed molecular docking, atomistic molecular dynamics (MD) simulations and free energy calculations to investigate the binding mode of the transition state analogue (TSA) inhibitor L-685,458^16^ (see Figure 1). The predicted binding pose of the inhibitor matches the binding geometry of C99 residues L49, V50, M51 and L52 and also mimics the beta-sheet formation observed for substrate binding near the active site of the enzyme.

**Figure 1:**
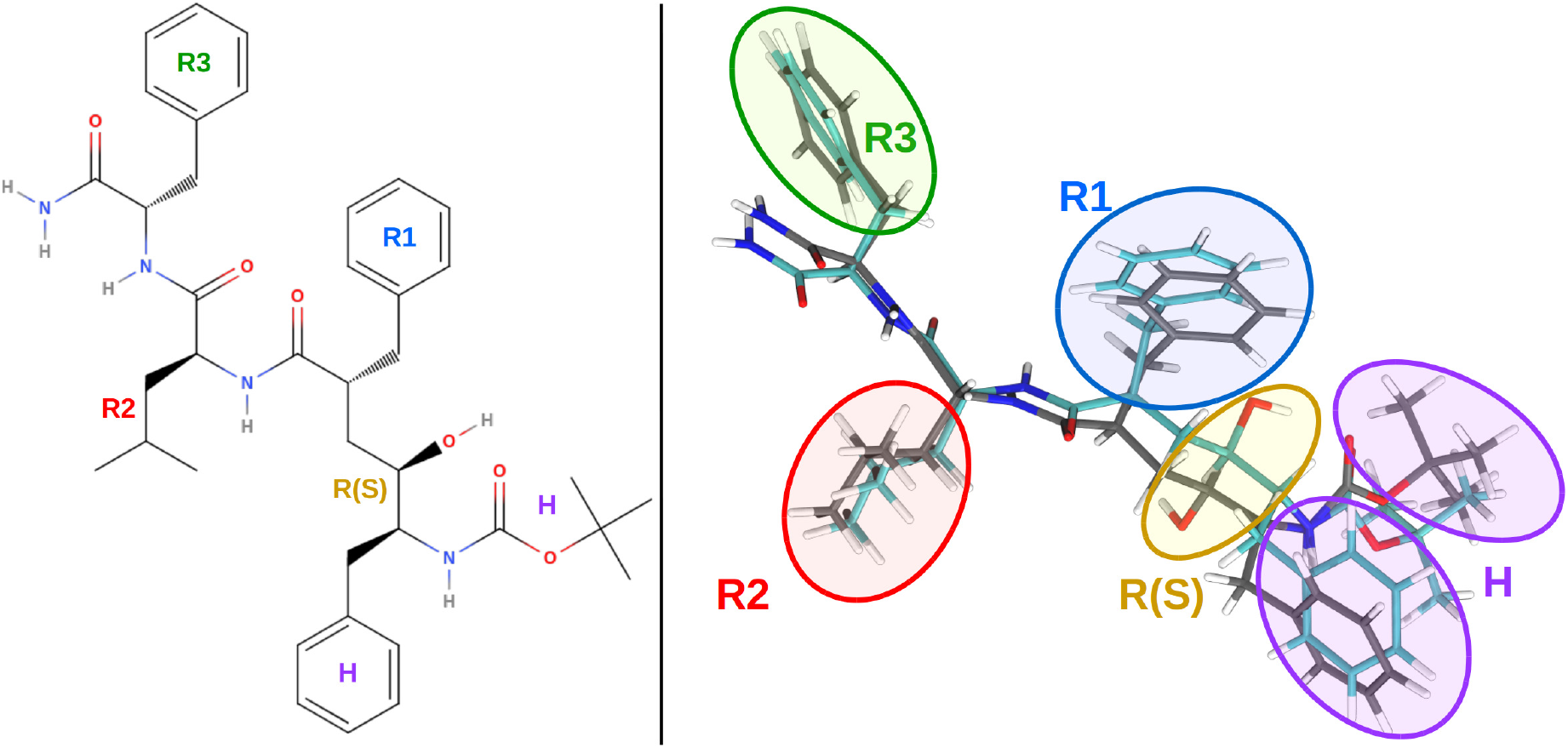
Structures of the TSA inhibitors L-685,458 and L-682,679. The three side chains occupying the proposed S1’, S2’ and S3’ pockets are termed “R1”, “R2”, “R3”, respectively. The other two large structural moieties of the ligand, pointing towards PS TMDs 2 and 3 in the bound structure, are denoted as “H” (”head group”). Left panel: Skeletal formula of the TSA inhibitor L-685,458. Right panel: Superposition of the optimized geometries of L-685,458 (grey) and L-682,679 (cyan).

We investigated all four possible presenilin active site (D257 and D385) protonation states and found that L-685,458 binds stably to three of them. The binding affinity, however, decreased with increasing residual charge on the active site residues, indicating pH sensitivity. The greatly reduced inhibitory performance^16^ of L-682,679, an epimer of L685,458 (see Figure 1 for comparison) can also be explained by our study.

Starting structures for inhibitors L-685,458 and L-682,679 were generated by geometry optimization (on B3LYP^36^ level, with the TZVP^37^ basis set, see Supplementary Information). Alignment of L-685,458 to GSEC-bound substrate residues V50, M51 and L52 (coordinates taken from a short simulation based on PDB structure 6IYC^24^) resulted in a placement of the inhibitor’s OH group, coinciding with the carbonyl oxygen of the scissile peptide bond in the GSEC-substrate complex. Additionally, the R1-R3 side chains of the inhibitor pointed in the same direction as the corresponding side chains of the APP fragment (V50, M51 and L52), hence also reproducing the down-up-down sidechain sequence of the enzyme-bound substrate. This placement, exactly mimicking the local substrate – enzyme interactions, appeared to be unique since in extensive docking searches no other sterically feasible placement could be identified with the OH group in a state of mimicking a transition state scenario.

To investigate the dynamics and stability of the docked geometry, MD simulations (1000 ns, including a POPC membrane and water) were performed for all four possible active site protonation states.

Creating starting structures for the S-epimer proved to be more challenging, because only rotamers different from the minimzed structure could fit in similar fashion as L-685,458. We selected two different ligand rotamers which were able to form many ligand-enzyme interactions for simulation: One, that was closer to the energy-minimum structure of L-682,679 (”P1”) and an alternative one that was more similar to the L-685,485-GSEC starting structure (”P2”) (Supporting Information, Figure 1). For each pose we generated simulation trajectories of 500ns, totaling 1000ns for every investigated PS protonation state.

The simulations where both catalytic aspartates were protonated are referred to as either “R-DP” (L685,458) or “S-DP” (L682,679). Simulations with a single protonated active site are termed “R-D257” and “S-D257” (protonated D257) or “R-D385” and “S-D385” (protonated D385), respectively. The states with non-protonated active site residues are denoted as “R-NP” or “S-NP”.

In simulations with at least one protonated catalytic residue, inhibitor L-685,458 maintains a stable hydrogen bonding network with PS residues G382, K380 and L432 throughout nearly all of the accumulated 3*μ*s of simulation time. Additionally, the OH group situated at the epimeric center binds strongly to the catalytic residues of GSEC (see Figure 2 for details). Since the main part of the ligand is situated exactly where Notch and C83 (C99) form beta-sheets with the enzyme,^23, 24^ only the head group protrudes into the binding site of the substrate helices. This explains why it is possible to inhibit GSEC without simultaneously preventing substrate binding.^14, 15, 39^

**Figure 2:**
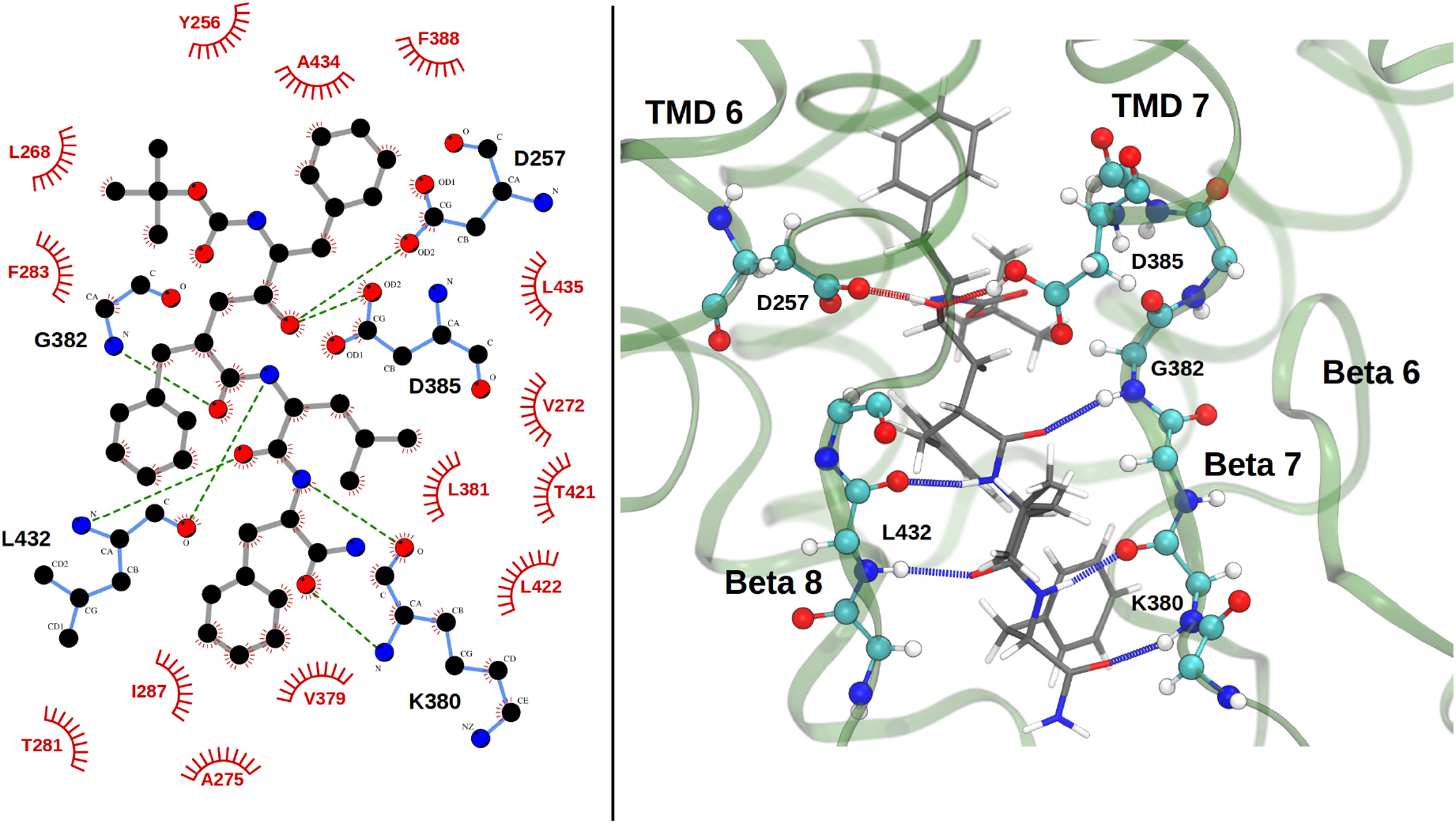
GSEC – L-685,458 interactions. Left panel: Interaction diagram (adapted from LigPlot+^38^ output). The green lines depict hydrogen bonds between the enyzme (blue bonds) and the ligand (grey bonds). Residues involved in hydrophobic interactions are shown in red. Right panel: Representative snapshot from simulation R-D385. Hydrogen bond colors correspond to the type of the donor atom, GSEC backbone is depicted in green. Carbons of GSEC residues are colored in cyan, those of L-685,458 are grey.

Figures 3 a) and b) illustrate that L-685,458 very closely resembles the transition state of the substrate. Superposition of L-685,468 (from simulation R-D385) and the GSEC-bound APP fragment (taken from a short simulation of the complex between the natural substrate and the WT enzyme in the same protonation state), shows that the TSA occupies exactly the same binding cavities as the substrate, explaining the high affinity and inhibitory efficiency of L-685,458.

**Figure 3:**
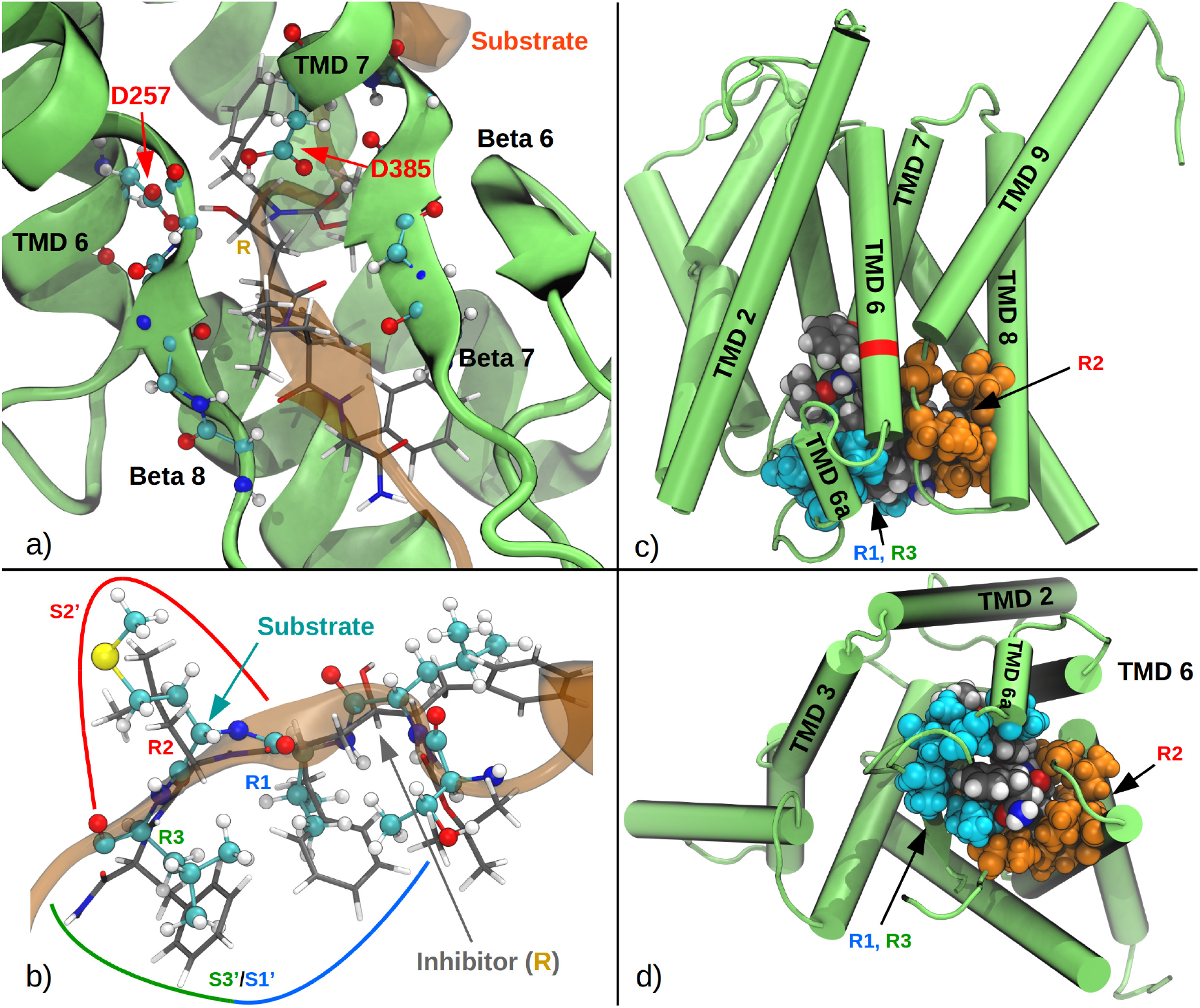
(a) Superposition of GSEC structures containing either the natural substrate or L-685,458 (a rotated close up view shown in (b)). Superposition involves the backbone atoms of the substrate binding site of PS. GSEC (green) and APP (orange) structures shown are from a short simulation of the native enzyme-substrate complex. The inhibitor (gray) is taken from simulation R-D385. (c,d) Location of the side chain binding pockets S1’, S3’ (occupied by R1 and R3, blue) and S2’ (occupied by R2, orange).

The structure of the bound inhibitor also revealed the location and constitution of the S1’, S2’ and S3’ binding pockets in GSEC, proposed by Wolfe et al.^4^ Interestingly, instead of three separated pockets, we identified only two: One S2’ pocket, which is very confined and situated close to TMDs 8 and 9 (see also Figure 3c and d). This pocket is permanently occupied by the smaller side chain R2. The putative binding pockets S1’ and S2’, on the other hand are part of the same spacious, hydrophobic cavity in which side chains R1 and R2 bind simultaneously (cf. with Figure 3c and d). These two finding offer an explanation for experimental results according to which the S2’ pocket is very sensitive to the size of the side chain, while S1’ and S3’ are not.^4, 40^

Table 1 provides a statistical overview over the nature of the residues constituting pockets S1’, S2’ and S3’.

**Table 1:** Interaction frequencies (in percent of sampled frames) between PS residues and R1-R3 of the bound L-685-458 inhibitor as well as the head group of the ligand (data taken from simulation R-D385). Only amino acids that are within 5Å of one of the side chains in at least 25% of sampled frames are shown here. *) Sites of known FAD mutations. †) only the residue’s backbone interacts with respective inhibitor side chain.

| R1, R3 |  | R2 |  | Head |  |
| --- | --- | --- | --- | --- | --- |
| K380 | 99% | L381* | 100% | L268 | 96% |
| V272* | 98% | L425 | 100% | I253 | 95% |
| I287 | 96% | T421 | 100% | F388* | 93% |
| T281 | 91% | A434* | 100% | G384*† | 88% |
| L282* | 89% | L422 | 99% | I143* | 83% |
| L271* | 84% | K380† | 95% | L435* | 70% |
| A275* | 73% | V379 | 93% | Y256* | 61% |
| L268 | 66% | L432 | 92% | F283 | 47% |
| F283* | 56% |  |  | L150* | 43% |
| L381*† | 27% |  |  | D257 | 38% |
|  |  |  |  | M146* | 29% |

In simulations featuring L-685,458 only R-NP exhibited permanent, partial dissociation of the ligand. Here, the hydrogen bonds to G382 and K380 were disrupted after just 80ns of simulation time. This is likely due to electrostatic repulsion of D257 and D385 (both are negatively charged in the R-NP simulation), thereby tearing apart the beta sheet and attracting a larger number of water molecules into the binding site.

According to experimental measurements,^16^ the affinity of GSEC for L-682,679 is much lower than for L-685,458 (IC_50_ value of >10000 nM, vs 17 ± 8 nM). In accordance with experiment, also in our simulations the enzyme-ligand stability was greatly reduced in the L-682,679-GSEC complex: In all simulations, partial dissociation events took place. Most frequently the beta-sheet was disrupted but also dissociation from the active site aspartates took place in more than one case (Supporting Information, Figure 2 shows a representative snapshot of L-682,679 binding).

The differences in affinity of the two epimers is also reflected by molecular mechanics generalized born surface area (MMGBSA) calculations, conducted on either the first 100ns (500 frames) of each simulation or the complete 1000ns trajectories (500 frames): The binding energies of the L-685,458 (R-epimer)-GSEC complexes are consistently more favorable for all simulations where at least one of the aspartates was protonated. The results are given in Table 2 and indicate that the R-DP simulation features by far the most stable enzyme-ligand complex if the complete trajectories are taken into account. Due to the large system size, no conformational entropy change estimations have been applied, therefore absolute numbers are not comparable to experiment. However, the predicted relative binding free energy difference between L-685,458 and L-682,679 amounts to 5-10 kcal/mol and agrees qualitatively well with experiment (1000 fold difference in Kd corresponds to a binding free energy difference of 4.2 kcal/mol)

**Table 2:** Summary of MMGBSA results, average water hydration numbers and ligand RMSFs. MMGBSA results are given for the first 100ns of each simulation (first results column) and all available simulation frames (second results column). Errors are given as standard errors (MMGBSA) or standard deviation (H_2_O and RMSF) *) P1 und P2 trajectories have been merged for evaluation.

| System | $\Delta G(100\text{ns})$ | $\Delta G(1\mu\text{s})$ | Mean # H<sub>2</sub>O | RMSF in Å |
| --- | --- | --- | --- | --- |
| **R-DP** | $-80.61 \pm 0.21$ | $-82.13 \pm 0.20$ | $7.76 \pm 2.22$ | 0.89 |
| **R-D257** | $-81.77 \pm 6.16$ | $-67.74 \pm 0.56$ | $10.90 \pm 3.30$ | 1.48 |
| **R-D385** | $-79.74 \pm 0.20$ | $-74.24 \pm 0.27$ | $11.88 \pm 2.96$ | 1.44 |
| **R-NP** | $-74.31 \pm 0.36$ | $-55.51 \pm 0.35$ | $31.25 \pm 6.42$ | 1.72 |
| **S-DP P1** | $-69.39 \pm 0.21$ | $-64.12 \pm 0.30^*$ | $16.83 \pm 5.04^*$ | 2.04* |
| **S-DP P2** | $-57.08 \pm 0.33$ | | | |
| **S-D257 P1** | $-60.22 \pm 0.24$ | $-58.53 \pm 0.32^*$ | $22.37 \pm 7.38^*$ | 2.45* |
| **S-D257 P2** | $-44.50 \pm 0.47$ | | | |
| **S-D385 P1** | $-74.02 \pm 0.24$ | $-63.49 \pm 0.35^*$ | $18.47 \pm 4.47^*$ | 1.91* |
| **S-D385 P2** | $-73.11 \pm 0.19$ | | | |
| **S-NP P1** | $-58.51 \pm 0.25$ | $-58.27 \pm 0.33^*$ | $22.27 \pm 7.25^*$ | 2.00* |
| **S-NP P2** | $-61.45 \pm 0.44$ | | | |

Furthermore, we calculated the mean number of water molecules within 4Å of the ligand to estimate the level of desolvation. The results show, that there is a large gap in solvent accessibility between the high affinity (R-DP, R-D257, R-385) and the low affinity (R-NP, S-DP, S-D257, S-D385, S-NP) complexes (table 2), with the former being significantly more desolvated.

Calculation of the root mean square fluctuations of the ligand painted the same picture: Simulations of the high affinity complexes, featured smaller ligand fluctuations, indicating more stable binding (ligand RMSD plots for each simulation are given in the Supporting Information, Figure 3). In summary, especially at pH values below pH 7 L-685,458 is expected to bind very stable to PS, closely mimicking the substrate, while the predicted affinity of L-682,679 is greatly reduced (in agreement with experiment).

Binding of the inhibitors to the enzyme active site involves slight conformational changes of the inhibitors to accommodate to the binding regions. In order to estimate this contribution we optimized the geometries of both ligands (bound state) to the nearest minimum (B3LYP/TZVP-level in vacuo). We found that the energy difference is higher for the low affinity inhibitor L-682,679 (97.89 kcal/mol) compared to L-685,458 (91.39 kcal/mol). This contribution further disfavors binding of L-682,679 and explains the observed tendency to perturb the binding region and to initiate complex dissociation.

Based on the identified binding mode it is now possible to make predictions regarding the possible improvement of inhibitory efficiency: For example, the amide tail of bound L-685,458 is close to the polar side chains of PS residues K380 and Q276 (minimum distances of approx. 2.8 and 3 Å in simulation R-D385, respectively). Hence, a hydrogen bond acceptor instead of the NH2 group at this position could further increase affinity and specificity of the inhibitor. Another potential improvement is enabled by the proximity between T421 and the R2 side chain: If adapted to contain a hydrogen bond acceptor R2 might be able to form a hydrogen bond with T421.

In conclusion, the ligand binding mode predicted by this study is in excellent agreement with available experimental data and can serve as a starting point to systematically explore possible GSEC ligands and to rationally modify existing inhibitors.

## Supporting Information

Materials and Methods section, Figures 1-3. For further reference we also provide a PDB file containing the structure of the predicted *γ*-secretase – L685,458 complex.

## Author Informations

M.H. performed research, analyzed data, and wrote the article; and M.Z. designed research and wrote the article.

## Acknowledgments

Financial support by the DFG (German Research Foundation) grant FOR 2290 (project P7) is gratefully acknowledged. Computer resources for this project have been provided by the Gauss Centre for Supercomputing/Leibniz Supercomputing Centre under grant pr27za.

