## Supplemental Information for "Uncovering the Binding Mode of *γ*-Secretase Inhibitors"

### Uncovering the Binding Mode of $\gamma$-Secretase Inhibitors: Supplemental Material

*M. Hitzenberger and M. Zacharias*

Physics Department T38

Technical University of Munich

James-Frank-Str. 1

85748 Garching, Germany

### Materials and Methods

#### Molecular Dynamics Simulation

All MD simulations were performed utilizing the GPU accelerated CUDA[1] version of PMEMD[2–4] that is part of the AMBER18[5] package. The proteins and POPC lipids were described by the AMBER14SB[6] and Lipid14[7] force fields, respectively. The TIP3P[8] water model was used to represent the aqueous environment. The inhibitor molecules were parametrized using the Antechamer[9] module of AMBER18, utilizing Amber's GAFF[10] force field. Prior to assigning RESP partial charges[11,12] (Hartree-Fock[13–16] at 6-31G* level[17–21]) to the ligands, the structures were optimized to a gas-phase energy minimum, using B3LYP[22] and the TZVP basis set.[23] All QM calculations have been performed with GAUSSIAN09.[24]
In the simulations, all pairwise non-bonded interactions were calculated up to distances of 9Å. Long-range electrostatic interactions (beyond 9Å) were calculated by the particle mesh ewald (PME) method.[25] Periodic boundary conditions were applied with box dimensions of approx. 107Å by 108Å by 211Å, containing besides GSEC and ligands, approx. 60300 water, 0.15M KCl as well as 304 POPC molecules.
The target temperature was set to 303.15K by applying the Langevin thermostat[26] with a collision frequency of $1ps^{-1}$. To achieve an NpT ensemble, the Monte Carlo barostat[27] was employed to maintain a constant pressure of 1.0 bar. In order to allow for time steps of 4.0fs, the SHAKE algorithm[28] was used, alongside with hydrogen mass repartitioning of the solute.[29]
Data analysis was carried out using CPPTRAJ[30] and VMD.[31] The latter was also used for rendering screenshots of the simulations.

#### Simulation System Setup

All simulations were based on the PDB structure 6IYC[32] representing GSEC in complex with the C-terminal fragment of the amyloid precursor protein (APP). However, the Cryo-EM structure missed large parts of the long, intracellular loop 6, a section of the N-terminal loop region of the substrate and more importantly also the position of the side chain of D385 (due to originally being mutated to Ala[32]). In addition, the carbonyl oxygen of L49 was not in contact with D257. Since both ligands are transition state analogues we performed a short MD simulation of the restored WT enzyme-substrate complex (in silico mutation of A385 to D385) to find a conformation in which the substrate is close to being cleaved (with both, D257 and D385 in close contact with the substrate as well as with each other[33]).
To achieve this, we first uploaded the PDB file to the OPM server[34] to calculate the orientation of the protein complex in the bilayer. Next, the re-oriented structure was embedded in a POPC bilayer and water by using the CHARMM-GUI server.[35] All missing (non-terminal) residues have been restored with the help of CHARMM-GUI and the modeled loop 6 was cleaved between residues M298 and A299 to represent the matured GSEC complex.[36]

Since a large section of the N-terminus of nicastrin was missing, we capped the structure with an acetyl (ACE) cap. An additional ACE group was added to presenilin. The C-termini of Aph-1a and the substrate were capped with N-methylamide (NME).
The protonation states of the titratable groups were assigned by using the PDB2PQR server[37] (pH was set to 6.5). Only the protonation state of the active site was chosen manually (two simulations, with either D257 or D385 protonated have been performed). The prepared structure was then relaxed, heated and pressurized in a 7-step equilibration procedure (see details on Table 1).
During initial equilibration, positional restraints on the solute were used to allow equilibration of the lipid and aqueous environment. Production runs have been performed without the use of any restraints.

| | **Steps** | **Time Step** | **K Amino Acids** | **K Lipids** | **Temperature** | **Pressure** |
|---|---|---|---|---|---|---|
| **min** | 50000 | minimization | 10.0* | 2.5 | — | — |
| **eq 1** | 25000 | 1fs | 10.0 | 2.5 | 0K to 303.15K | NVT |
| **eq 2** | 25000 | 1fs | 5.0 | 2.5 | 303.15K | NVT |
| **eq 3** | 25000 | 1fs | 2.5 | 1.0 | 303.15K | 0bar to 1bar |
| **eq 4** | 50000 | 2fs | 1.0 | 0.5 | 303.15K | 1bar |
| **eq 5** | 50000 | 2fs | 0.5 | 0.1 | 303.15K | 1bar |
| **eq 6** | 50000 | 2fs | 0.1 | 0.0 | 303.15K | 1bar |

**Table 1:** Overview over the 7 equilibration steps performed for all simulations. Positional restraints, K are given in kcal/mol*Å$^2$. The functional form of the restraints is: K*$\Delta x^2$.
*) For simulations with a docked inhibitor, no restraints were applied to the protein or the inhibitors during initial minimization.

##### Generation of Starting Structures of GSEC-Inhibitor Complexes.

The substrate-GSEC complex structure was equilibrated during a 300 ns unrestrained MD simulation at constant temperature of 303.15 K and constant pressure of 1 bar. The equilibrated structure (with protonated D257) was then used as a template for the generation of the initial enzyme-inhibitor structure. Docking of the transition state analogue inhibitor L-685,458 was achieved by superposition onto selected residues of the APP fragment near the active site. The superposition of the inhibitor onto substrate residues 49 to 52 (backbone atoms) resulted in a structure with the inhibitor occupying exactly the same cavities as the corresponding side chains of residues 49 to 52 of the APP fragment (with basically no sterical overlap) and was able to form the same hydrogen bond interactions as the corresponding patch of the natural substrate. In addition, it placed the OH group of the inhibitor into the vicinity of the active site aspartate residues in order to form the transition state inhibition.
After short energy minimization this initial conformation was used for all L-685,485 simulations with different active site protonation states. The enzyme, bilayer and water coordinates were carried over from the GSEC-substrate simulation to form the start structure. Prior to the 1000ns unrestrained production runs, the equilibration procedure shown on table 1 was

performed.
In case of the second investigated ligand it was necessary to create several rotamers in order to get the OH group to interact with the catalytic side chains while ensuring the formation of the beta-sheet between ligand and protein. The two most promising structures have then been selected for simulation (see Figure 1). To ensure the same amount of sampled complex frames for both ligands, after equilibration (Table 1) 500ns trajectories have been generated for each pose and protonation state.

#### Evaluation of sampled trajectories

For the estimation of enzyme ligand affinities, the post-processing single-trajectory molecular mechanics Generalized Born molecular mechanics (MMGBSA) method[38,39] (as implemented in the MMPBSA.py Amber-module[40]) has been utilized. Since the area in which the ligands were binding to GSEC was accessible to water, the permittivity of the outside region was set to 78.3, while the internal permittivity of the protein was assumed to be 1.0. To stay consistent with the simulation, the salt concentration was set to 0.15M. To calculate the nonpolar part of the sovlation free energy, the surface tension was set to 0.0072 kcal/mol/$Å^2$.

To compute the root mean square fluctuations, RMSF of the ligand (given as mean of all ligand atoms in the main manuscript), all frames of the respective simulations have been aligned according to the backbone atoms of presenilin (excluding the highly mobile loop 6 region). Subsequently, the fluctuations of the ligand were calculated.

#### Supplemental Figures

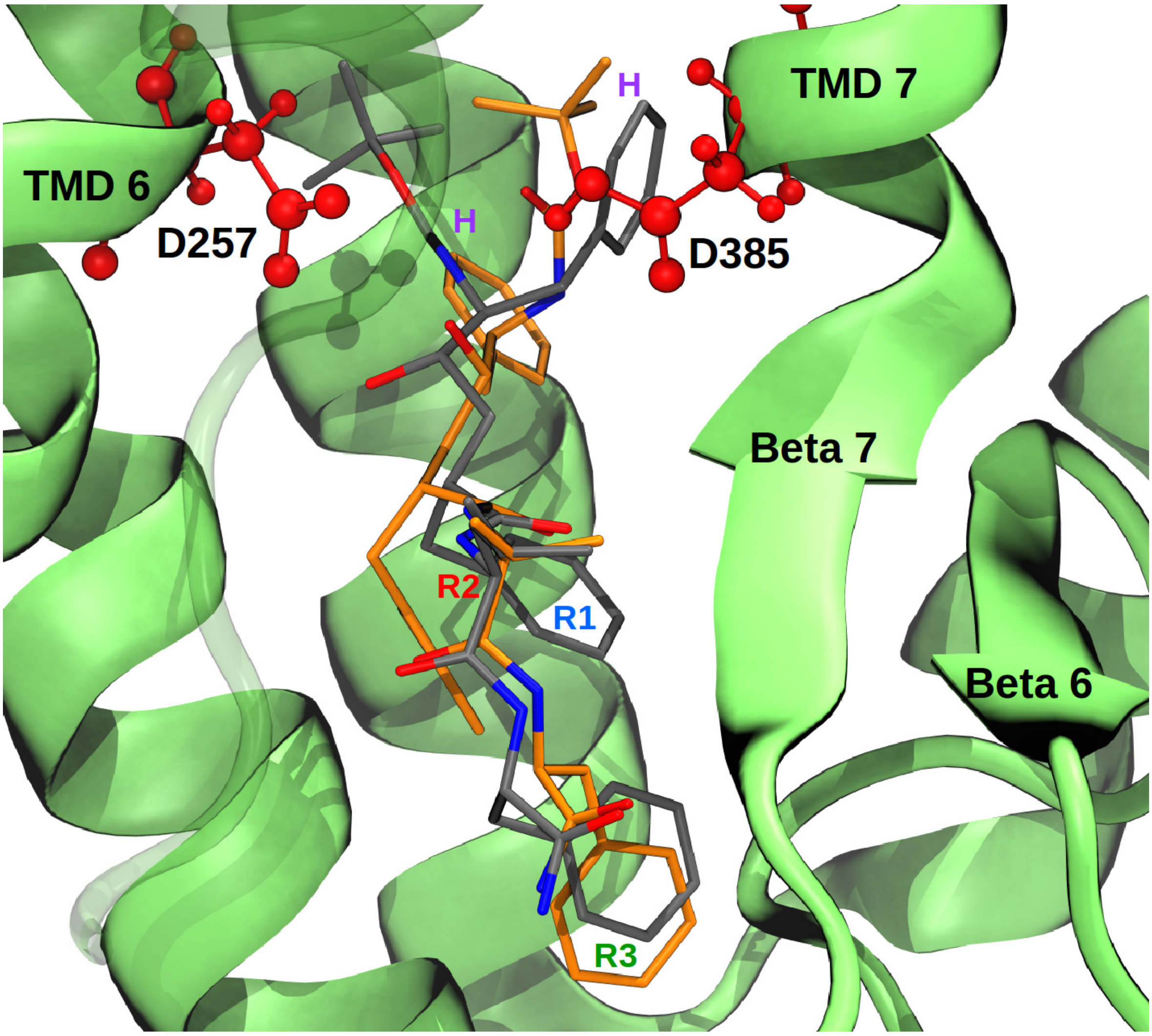


**Figure 1:** The two different initial L-682,679 poses that have been selected for simulation: Pose 1 (P1) is depicted in grey, while pose 2 (P2) is shown in orange.

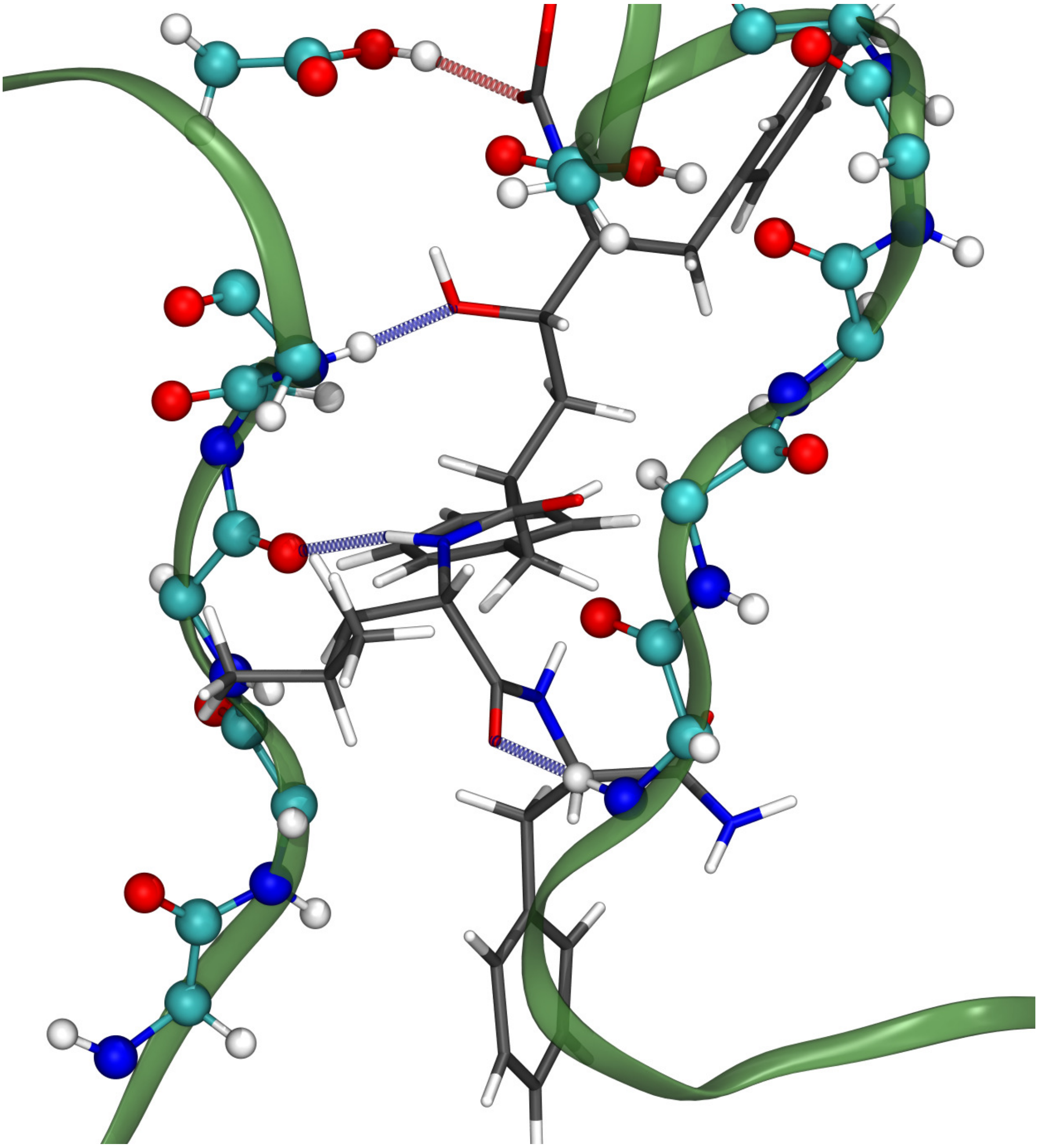

**Figure 2:** Representative L-682,679 binding snapshot, taken from simulation S-DP (P2). The protein backbone is depicted in green, while the ligand is colored in grey. The color of the hydrogen bonds coincide with the nature of the donor atom. Note, that the OH group dissociated from the active site and most of the beta-sheet hydrogen bond network has been disrupted as well.

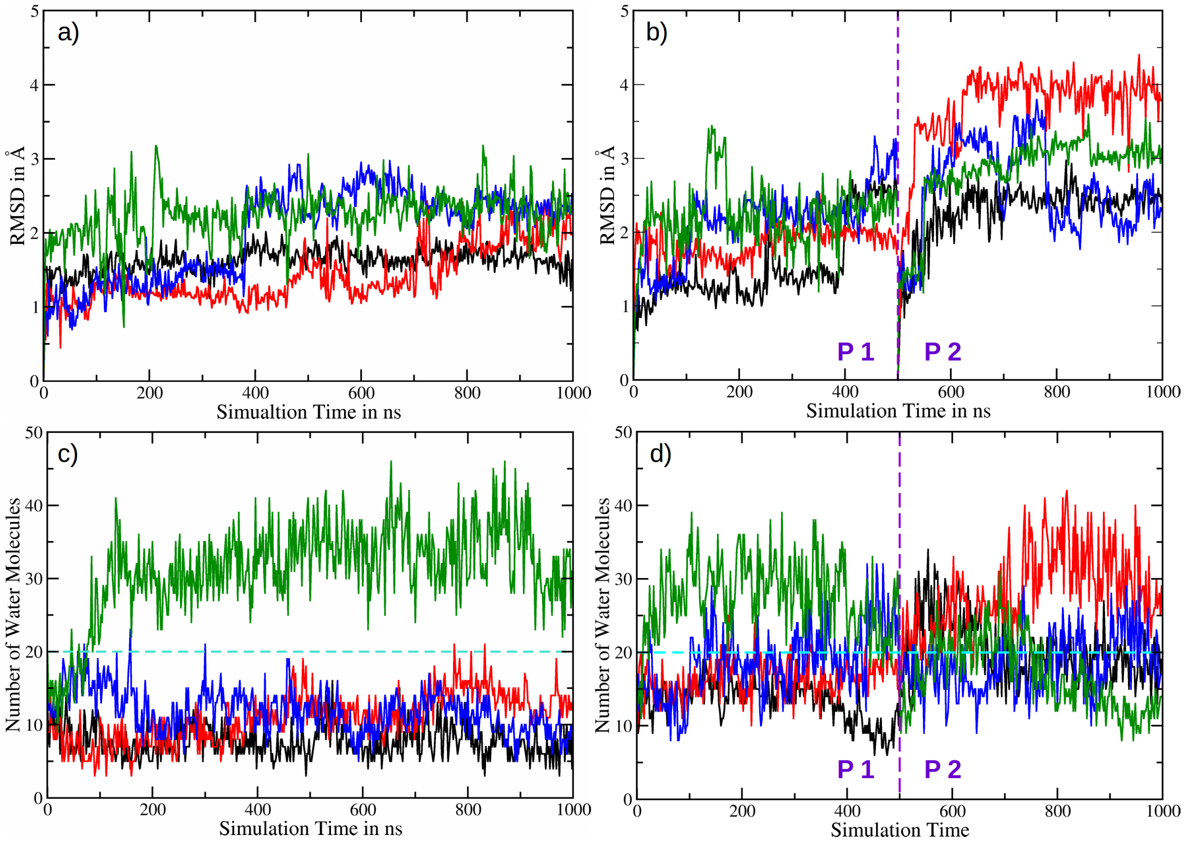


**Figure 3:** a) Ligand heavy-atom RMSDs of simulations R-DP (black), R-D257 (red), R-D385 (blue) and R-NP (green). The large change in RMSD around 400ns in simulation R-D385 is due to a re-arrangement of the head groups. The initial (slightly mis-aligned) orientation was stabilized by the deprotonated D257 side chain. After the re-arrangement, the tert-butyl carbamate group exactly imitated the orientation of the backbone of the natural substrate. In L-685,458-GSEC simulations with protonated D257 this re-arrangement happened on a much faster timescale and is therefore not captured by the RMSD calculations.
b) Ligand heavy-atom RMSDs of simulations L-DP (black), L-D257 (red), L-D385 (blue) and L-NP (green). The simulations of pose 1 (P1) are on the left of the divider (dashed, magenta line), while the P2 simulations are on the right. The first frame of each simulation has been used as reference.
c), d) Plots of water hydration. In every evaluation frame all water molecules within 4Å of the ligand were counted. Color coding is the same as above. For easier comparison a vertical line was added, denoting 20 water molecules.
